# Polygraph: A Software Framework for the Systematic Assessment of Synthetic Regulatory DNA Elements

**DOI:** 10.1101/2023.11.27.568764

**Authors:** Avantika Lal, Laura Gunsalus, Anay Gupta, Tommaso Biancalani, Gokcen Eraslan

## Abstract

The design of regulatory elements is pivotal in gene and cell therapy, where DNA sequences are engineered to drive elevated and cell-type specific expression. However, the systematic assessment of synthetic DNA sequences without robust metrics and easy-to-use software remains challenging. Here, we introduce Polygraph, a Python framework that evaluates synthetic DNA elements, based on features like diversity, motif and k-mer composition, similarity to endogenous sequences, and screening with predictive and foundational models. Polygraph is the first instrument for assessing synthetic regulatory sequences, enabling faster progress in therapeutic interventions and improving our understanding of gene regulatory mechanisms.

## Background

Cis-regulatory DNA elements (CREs) are DNA sequences that drive and tune gene expression, in part through the selective binding of transcription factors or other regulatory molecules necessary to initiate and maintain transcription [1]. Their cell-type specific activity makes carefully designed CREs a promising avenue for synthetic biology and DNA-based nucleic acid therapeutics such as cell and gene therapy [2–6]. Recent deep learning models trained on high-throughput sequencing data integrate the complex grammar embedded in DNA and serve as powerful tools to predict the activity of native and synthetically generated regulatory elements [7–11]. However, evaluating and prioritizing designed sequences remains challenging without a comprehensive understanding of their properties and potential regulatory mechanisms.

Current DNA design methods range from classic optimization algorithms [12–14] to generative modeling approaches [15–19] aimed at driving desired expression profiles, transcription factor binding patterns, and other functional outcomes. These methods invite exciting open questions: What regulatory mechanisms do they exploit? Do they converge on the regulatory grammar observed in natural sequences or identify novel combinations of binding motifs? How can sequences be systematically selected from millions of computationally generated options? Sequence prioritization is currently limited by (1) the lack of integrated software to predict and analyze CRE performance across biological contexts; (2) the absence of defined metrics for comprehensive CRE evaluation, making it difficult to standardize assessments across studies and design methods; and (3) poor understanding of the biological underpinnings of designed sequences, including how CREs may behave differently across cell types and tissues.

Here, we present Polygraph, a software package that enables systematic evaluation of designed DNA sequences through sequence analysis, transcription factor motif composition analysis, predictive modeling, and language modeling. By facilitating the comparison of design algorithms and prioritization of CRE candidates, Polygraph provides a path toward robust and interpretable CRE engineering.

## Results

### The Polygraph package

Polygraph is a Python package that accepts DNA sequences of any length, analyzes their properties including their predicted activity, and compares them to user-defined reference sequences such as genomic regulatory elements. Polygraph also supports statistical significance testing across groups and predictions using custom models. In this paper, we demonstrate the use of this package on two datasets: yeast promoters designed with directed evolution [8] and first-order optimization method Ledidi [13], as well as human enhancers designed by Gosai et al. [11] where the authors employed FastSeqProp [12], simulated annealing [20], and AdaLead [14].

#### 1. Sequence composition analysis

Polygraph includes tools to evaluate sequence composition metrics like GC content, k-mer and gapped k-mer counts (abundance of nucleotide subsequences of length *k*), and edit distance to reference sequences. K-mer or gapped k-mer composition is used to cluster sequences and visualize them in principal component analysis (PCA) space. We apply a *sequence diversity* metric, defined as the average k-nearest neighbor (KNN) distance between a sequence and its neighbors from the same group [11], to quantify how similar designed sequences are to each other. We also offer a function to train support vector machine (SVM) classifiers to empirically test whether synthetic sequences can be discriminated from native DNA. Human regulatory regions exhibit elevated GC content [21], enriched motifs like GATA [22], CpG islands [22], and short repeats [23]. Evaluating sequence composition with our quantitative metrics could therefore provide insight into the novelty and “humanness” of computationally designed regulatory elements.

#### 2. Transcription factor binding motif analysis

Motif content reveals which combinations of transcription factors can bind to a DNA sequence. Polygraph uses FIMO [24] to scan sequences against the JASPAR motif databases [25] and describe their regulatory grammar. We count individual binding sites and compute the enrichment of motif pairs. Motif scanning reports start positions, orientations, and spacing, indicating if placed motifs are centered and if different methods pursue different syntactic strategies. We apply non-negative matrix factorization (NMF) to decompose the motif count matrix into common transcription factor programs shared across sequences [11]. Similar to k-mer analysis, we visualize and cluster sequences by motif count, compute euclidean and KNN distance between sequence groups, and train classifiers to differentiate native and synthetic DNA. Model-driven *in silico* mutagenesis scores the importance of each motif. Altogether, motif analysis may be used to uncover higher-order multi-factor regulatory rules exploited by different design approaches.

#### 3. Predictive modeling

Polygraph integrates trained neural network models to evaluate designed sequences on key properties like activity, specificity, and chromatin accessibility. We provide three pretrained open-source models for yeast and human prediction, with the option of integrating custom PyTorch models. Polygraph enables flexible and fast model prediction on generated sequences, cell-type specificity evaluation, and PCA or UMAP [26] visualization of sequence embeddings. These latent vector representations, created by evaluating sequences using the lower layers of the neural network model, may capture functionally relevant properties better than k-mer or motif representations alone. In addition, we provide a “guided evolution” function to evolve DNA sequences with high predicted activity while maintaining similarity to reference native sequences, based on their Euclidean distance in the model’s latent embedding space (see **Methods**).

#### 4. Language modeling

Autoregressive DNA language models are self-supervised models that aim to implicitly capture the rules of gene regulation as well as the rules of coding genes of the genome on which they are trained. These models can be queried to infer how likely a given sequence is sampled from the training set which in the case of human DNA language models is the human genome. Here we use HyenaDNA [27] to quantify the log-likelihood of synthetic sequences. These scores represent their “humanness” which is used as a proxy metric of how realistic generated sequences are.

### Yeast promoter analysis

We applied Polygraph to comprehensively analyze native and computationally designed 80 base-pair (bp) long yeast promoters. We consider 50 native yeast promoters with high reported expression in media and 50 with low reported expression [28,29]. We additionally generated 50 synthetic promoters using directed evolution [8] and 50 using gradient-based optimization with Ledidi [13], and 50 using Polygraph’s guided evolution function which maximizes predicted activity while maintaining similarity to native sequences (see **Methods**).

Using Polygraph, we ask how designed sequences might drive expression and if they resemble native yeast sequences. Promoters created using both evolution and gradient methods exhibited higher GC content than their random starting points, unlike native DNA (mean: 53%, 59%, 50%, respectively, evolution p-adj = 9.51e-02, gradient p-adj = 1.20e-05, Kruskal-Wallis, **Figure 2a, Supplementary Table 1**). This impact was more modest for sequences produced by guided evolution (mean: 47%, p-adj=9.51e-02), which encourages a resemblance to native sequences. K-mer analysis revealed that the designed promoters were enriched for GC-rich patterns like ‘CAGCC’ (gradient p-val = 5.26e-04, evolution p-val = 7.89e-04, guided evolution p-val = 5.26e-04, Kruskal-Wallis) and ‘GGCAC’ (gradient p-val = 5.26e-04, evolution p-val = 5.26e-04, guided evolution p-val = 7.89e-04, Kruskal-Wallis, **Supplementary Table 2**). Edit distances to native high promoters were comparable between designed and random sequences, suggesting that synthetic promoters did not grow to more closely resemble effective native sequences through iterative design (**Figure 2b**). Although designed sequences drive high predicted expression, they do not resemble the yeast genome.

**Figure 1:**
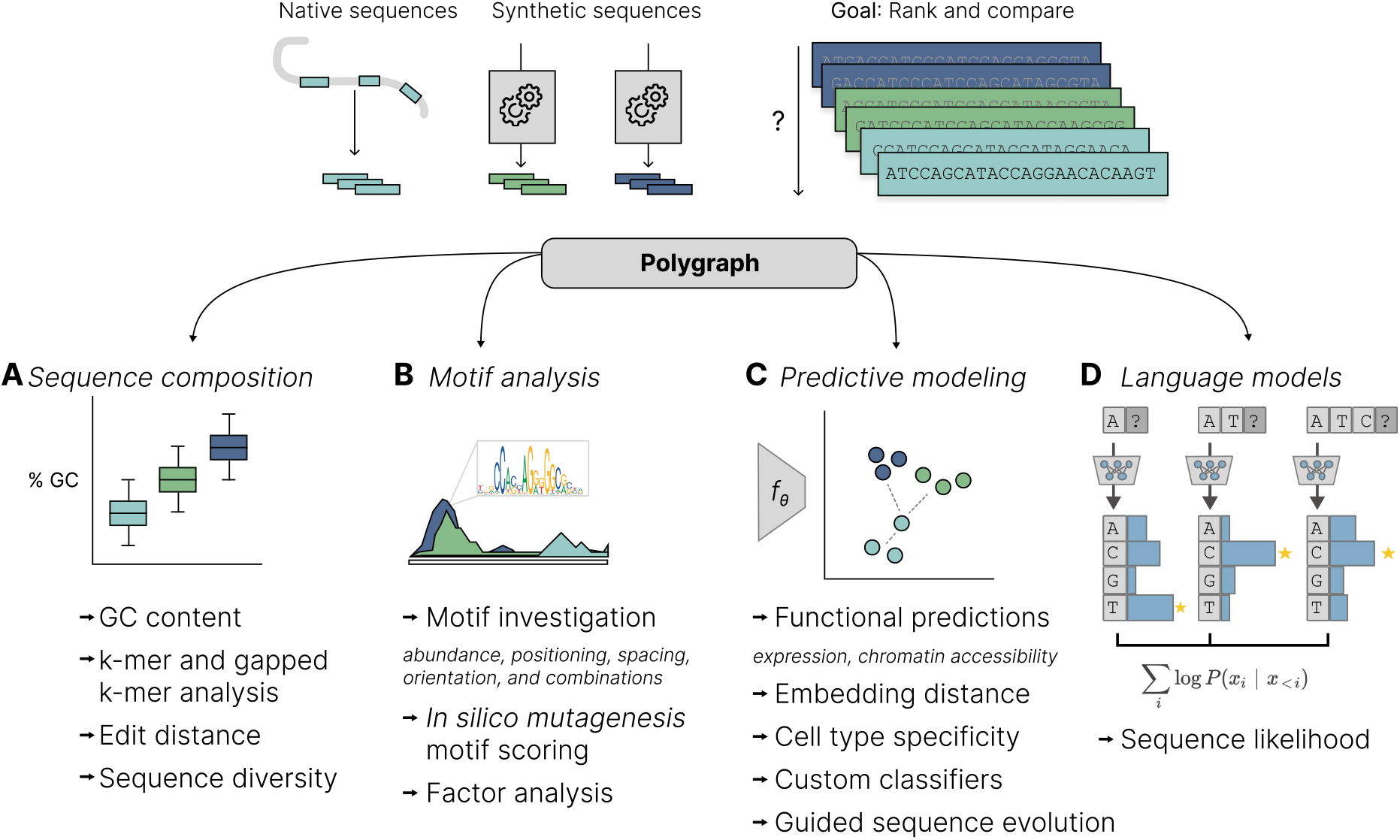
Schematic of Polygraph. Polygraph is a comprehensive package to evaluate and compare native and synthetic DNA sequences. Polygraph implements four classes of analysis: (A) Sequence composition, which evaluates GC content (percentage of bases in a sequence that are guanine (G) or cytosine (C), k-mer and gapped k-mer abundance, edit distance, and sequence diversity. (B) Motif analysis to quantify known transcription factor binding sites, analyze motif combinations, positioning, and importance, score motifs with in silico mutagenesis (ISM), and perform Non-Negative matrix factorization (NMF) to identify common transcription factor programs. (C) Predictive modeling to evaluate designed sequences based on key properties such as activity, specificity, and chromatin accessibility. The latent space obtained from intermediate layers of predictive models is used to visualize sequences and compute group diversity. These metrics can be used to design custom sequences with guided evolution. (D) Application of a human DNA language model to score native sequence likelihood.

**Figure 2:**
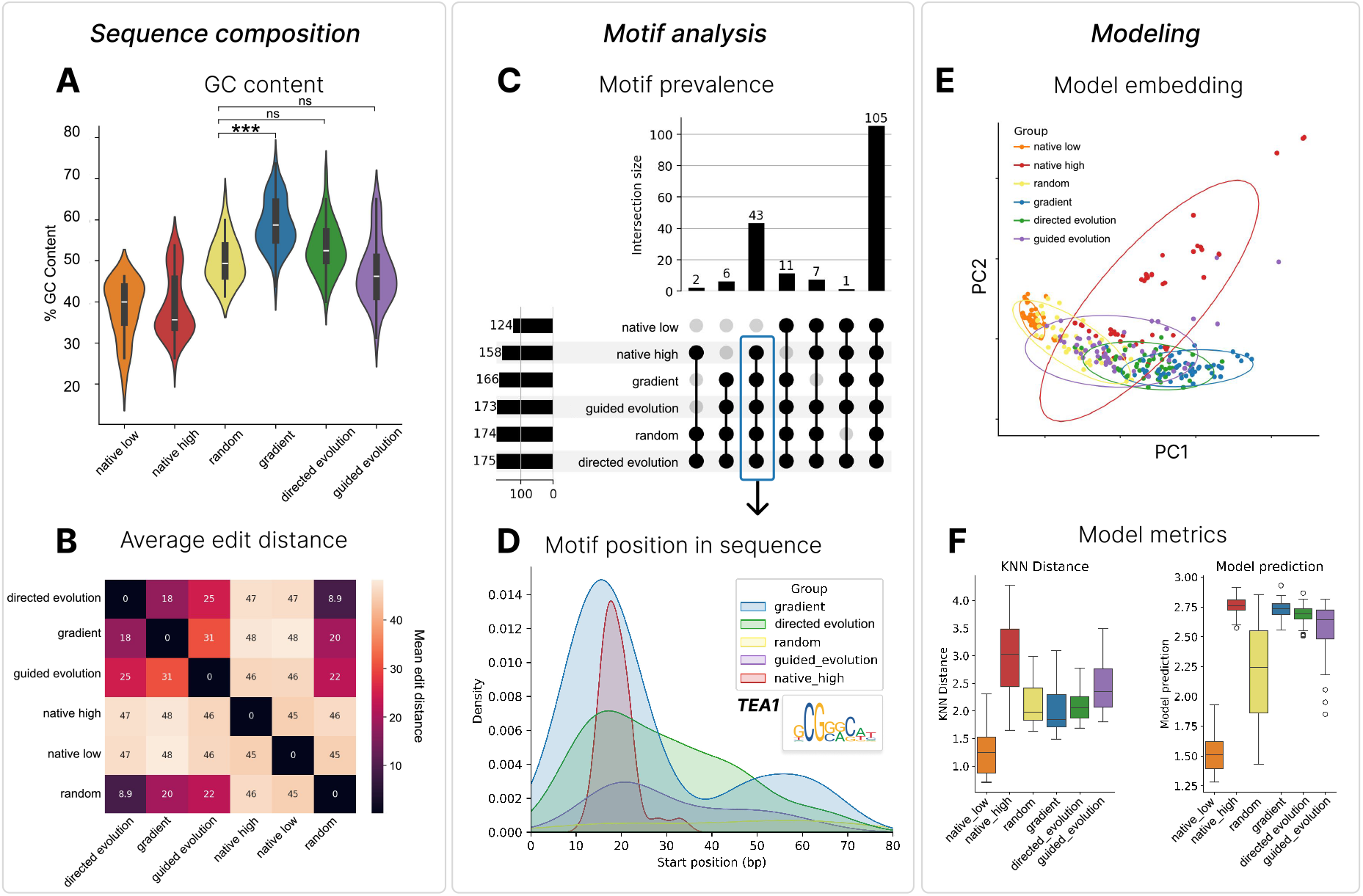
Sequence analysis of native, random, and designed yeast promoters. **a**. GC content distribution across native yeast promoters (labels: native_low and native_high), random sequences (random), and synthetic promoters designed via directed evolution (directed_evolution), gradient-based optimization (gradient), or guided evolution (guided_evolution). The native_high and native_low classes are native promoters with the highest and lowest measured expression, respectively. **b**. Average edit distance between sequences in all groups. **c**. Overlap of shared transcription factor binding motifs across native and generated promoter sequences. **d**. Distribution of TEA1 motif start locations, a motif absent in lowly expressed native promoters. **e**. PCA visualization of sequence embeddings from a sequence-to-expression predictive model. **f**. Nearest neighbor distance metrics in embedding space describe diversity within sequence groups (left). Predicted transcription from native, designed, and random sequences (right).

Next, we scanned against the 175 yeast motifs in JASPAR and identified 158 and 124 motifs in the native high and low groups, respectively (**Figure 2c**). 43 motifs were common to the highly-expressed native and all designed sets but absent in lowly-expressed native promoters. Conversely, 6 motifs appeared exclusively in synthetic sequences, which arose through the optimization process (CHA4, RDS2, STP2, SWI4, TBS1, cre-1; **Figure 2c, Supplementary Table 3**). We conclude that design methods can insert *de novo* motifs and that Polygraph can be a useful tool to identify them. Profiling the start position of shared motifs like TEA1 revealed common positioning, with binding sites frequently starting between positions 10-30, hinting at functional grammar (**Figure 2d**). This preference was stronger in Ledidi-generated sequences compared to directed evolution sequences, suggesting gradient-based methods may optimize motif arrangements more efficiently. Specific motif pairs like RSC30/RTG3 and RSC30/PDR3 were also enriched in designed groups (gradient p-adj = 2.75e-12, evolution pd-adj=3.76e-06, Kruskal-Wallis).

Finally, we evaluated the predicted transcriptional activity across sequences using an independent model trained to predict yeast expression from sequence. By inspecting, the sequence embeddings from intermediate layers of the neural network, we find that native sequences are the most diverse - native sequences have the most variance in embedding space, high intragroup KNN-distance, and high average distance to the closest sequence of the same type (**Methods, Figure 2e, f, Supplementary Figure 1**). As expected, native high promoters showed elevated predicted activity compared to native low promoters (**Figure 2f**). However, designed promoters reached similarly high predicted expression levels despite starting from random sequences, and one sequence generated by Ledidi was predicted to exceed the activity of all native promoters, demonstrating the ability of computational optimization to match or surpass the transcriptional activity of native regulatory DNA. In sum, these results point to a regulatory grammar underrepresented in native sequence, illustrating how integrating diverse *in silico* metrics provides insights into the sequence determinants underlying regulatory element design.

Interestingly, promoters generated by guided evolution were more similar to native high promoters than those generated by other methods while still achieving high predicted activity. No k-mers or motifs were differentially abundant compared to natives, whereas other methods showed significant differences (**Supplementary Table 2**). Guided evolution promoters had the least distance to the closest native sequence based on motifs (evolution p-val=0.108, gradient p-val=0.143, guided p-val=0.0824, **Supplementary Table 1, Figure 2e**). They were also most likely to be misclassified as native by k-mer or motif classifiers (**Supplementary Table 1**). Therefore, while existing tools tend to yield unnatural sequences, incorporating Polygraph metrics into design algorithms can produce effective sequences that mimic natural regulatory grammar.

### Human K562 enhancer analysis

We next applied Polygraph to evaluate 6,000 synthetic human enhancer sequences designed for the K562 cell line by AdaLead, FastSeqProp, and simulated annealing. These generative approaches were guided by an expression prediction model trained on a library of 776,474 genomic sequences tested in a massively parallel reporter assay (MPRA) for cell-line specific expression [11]. We compared these computationally designed enhancers to native human sequences from open chromatin regions specific to K562 cells and randomly generated DNA (**Methods**), again asking whether their composition and mechanisms resembled that of effective endogenous elements with high K562 specificity.

Sequence analysis revealed that while all sequences converge ∼50% GC content, designed sequences demonstrate lower GC variance (native = 49.2% ± 0.0655, simulated annealing = 50.0 ± 0.0260, **Supplementary Table 4**). K-mer distributions also differed markedly – ‘GGAG’, ‘CCAG’, ‘CCTG’ and ‘CAGG’ were common in native sequences, whereas ‘TATC’ and ‘GATA’ ranked as the top two k-mers across all design method (‘GATA’ p-val vs native sequences, FastSeqProp = 2.48e-295, AdaLead = 3.37e-233, simulated annealing = 1.71e-167, Kruskal-Wallis, **Figure 3a**). These designed sequences produced shared k-mer populations that differed not only from native sequence, but also from random starting sequence, suggesting a common shift toward more functional sequence grammar.

**Figure 3:**
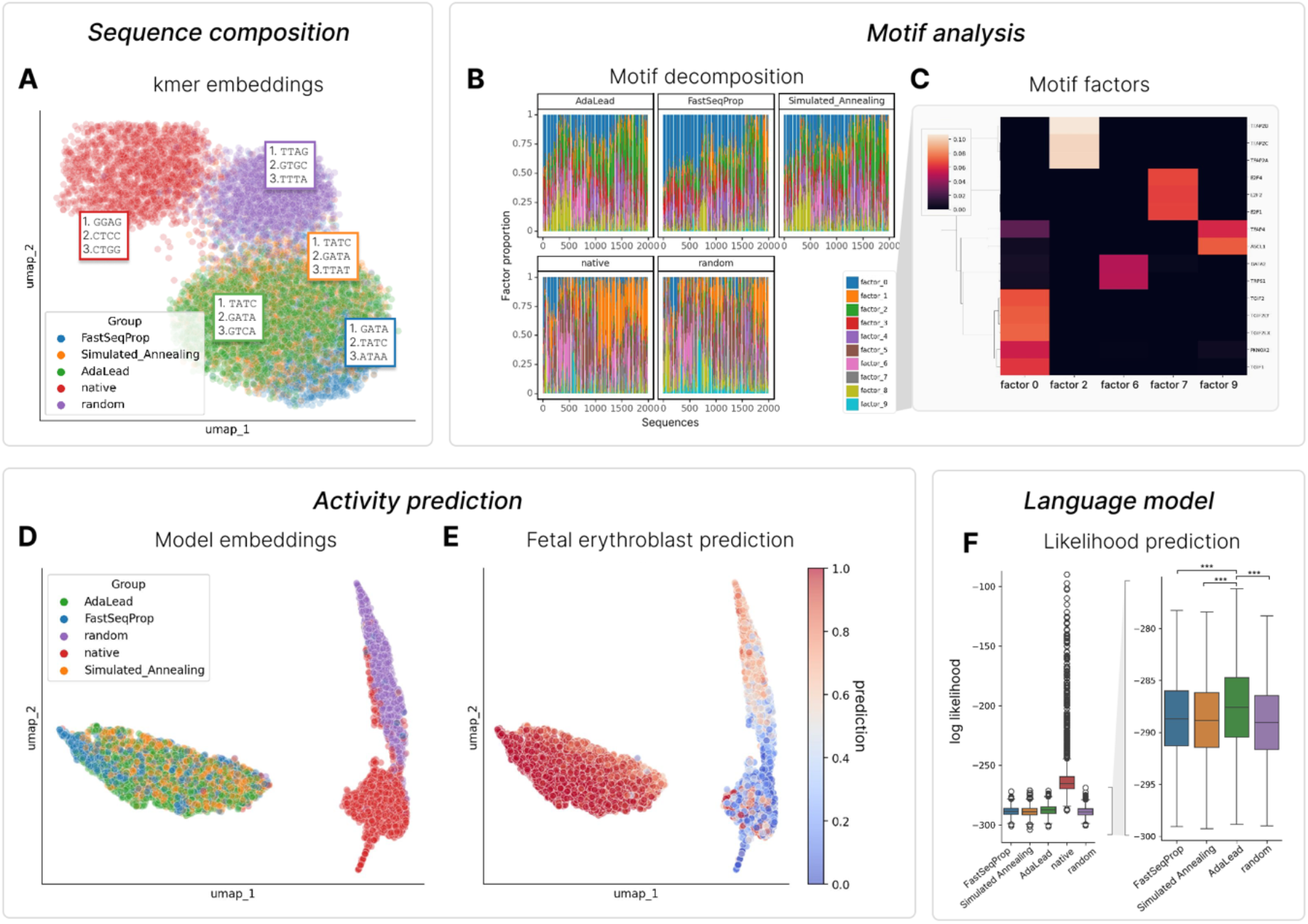
Sequence analysis of designed and native human enhancers in K562. **a**. UMAP visualization of 4-mer composition in uniformly random DNA, native DNA, and designed K562-specific enhancers generated with FastSeqProp, Simulated Annealing, and AdaLead. Top enriched k-mers are listed. **b**. Non-negative matrix factorization of motif counts, revealing distinct regulatory program preferences for designed and natural sequences. **c**. Motifs driving NMF components. **d**. Model embeddings from a chromatin accessibility model trained on fetal and adult human tissues[30]. **e**. UMAP of predicted chromatin accessibility in Fetal erythroblast, with designed sequences showing higher predicted erythroblast activity than native sequences. **f**. Likelihood predictions from a DNA language model, with only native sequencing scoring as probable human DNA.

Transcription factor binding motif analysis revealed substantial differences in regulatory grammar across groups. We scanned sequences for motif matches against 871 transcription factor motifs in the JASPAR database[25] (**Supplementary Table 5)**. NMF revealed that AdaLead, FastSeqProp, and simulated annealing produce highly similar compositions of motif factors, reliant on a diverse combination of motifs like TGIF1, TGIF2, PKNOX2 not observed in native or random sequences (e.g. average factor 0 loading z-scores = 0.42 (AdaLead), 0.55 (FastSeqProp), 0.31(simulated annealing), -1.46 (native), -2.16 (random), **Figure 3b,c, Supplementary**). In contrast, the ASCL1 motif featured prominently in native but not designed sequences. Sequence decomposition reveals how generative models converge on a shared vocabulary of motifs to produce synthetic K562 enhancers. Divergent motif strategies found in native and designed sequences can both produce cell-type specific regulatory sequences.

We next evaluate the activity of the K562-specific designed sequences in 30 adult and 15 fetal human tissues with 203 annotated cell types (**Supplementary Figure 3**). Using a binary sequence-to-chromatin accessibility model, we find that synthetic enhancers group together in embedding space, distinct from real or random elements aside from a small minority of native sequences (**Methods, Figure 3d**). Designed sequences are more diverse than both native and random elements, as measured by within-group KNN distance in embedding space (**Methods, Supplementary Table 4**). Open chromatin in fetal erythroblast, a blood cell type related to the K562 erythrocyte leukemia line, is predicted to be higher across synthetic enhancers than native sequences (mean prediction, AdaLead=0.95, simulated annealing=0.95, FastSeqProp=0.96, native=0.37, random=0.54, **Figure 3e, Supplementary Figure 4**), while predictions are low for the unrelated CD4+ T cell type. Designed sequences can identify sequence features that represent the preserved biology between cell lines and their origins in the human body.

Finally, we computed sequence likelihoods using an autoregressive DNA language model[27] by summing the predicted log-likelihood of each nucleotide across the sequence **(Methods**). As expected, native sequences showed high log likelihood **(Figure 3f)**. AdaLead-generated sequences had significantly higher log-likelihoods than FastSeqProp, simulated annealing, and random sequences (p-val = 1.54e-09, p-val = 4.85e-13, p-val = 7.66e-18, Kruskal-Wallis), suggesting greater similarity to genomic sequences.

## Discussion

The rational design of DNA elements holds great promise for both improving our understanding of gene regulatory mechanisms and driving cell-type specific expression in therapeutic contexts. While Polygraph provides an integrated toolkit for evaluating designed regulatory elements, several exciting directions remain to improve analysis and application. Integrating a wider array of predictive models, including those predicting chromatin accessibility, protein binding, histone modifications, and chromatin states will add more regulatory perspectives to sequence evaluation. Condensing our metrics to create a score with which to prioritize sequences will also be critical when screening millions of candidates for experimental validation. Experimentally validating a small number of synthetic sequences and iteratively feeding the results back to the generative design processes in an active learning framework is a promising future direction.

## Conclusions

As demonstrated in yeast and human case studies, Polygraph provides insights into the properties of computationally designed regulatory elements. Sequence analysis revealed how different generative algorithms converge on common grammar, often distinct from native DNA. Motif scanning uncovered shared transcription factor binding sites exploited for specialized activity like cell-type specific expression across methods. Predictive neural networks allow benchmarking design methods and resolving commonalities and differences between methods in terms of high-level regulatory sequence features that are implicitly parsed out by predictive models in an untargeted manner. DNA language modeling provides a “humanness” score describing similarity to genomic sequences. Incorporating these similarity metrics into “guided evolution” yielded synthetic regulatory elements predicted to have high activity while reducing extreme divergence from natural sequences.

Together, Polygraph sequence evaluations enable comprehensive assessment and comparison of regulatory element design approaches. This work establishes a robust computational platform to understand the regulatory code and advance the engineering of precise, tunable cis-regulatory elements for therapeutic applications going forward.

## Methods

### Polygraph package details

#### Sequence composition analysis

Polygraph calculates the following metrics of sequence composition for each provided sequence: Sequence length, GC content, edit distance with respect to reference sequences, k-mer frequency, gapped k-mer frequency, unique k-mers, and k-mer positions. Embedding metrics (described below) can be computed for each group of sequences based on their k-mer frequency. It uses statistical tests (described below) to compare the distribution of any of these metrics across groups.

#### Motif analysis

Polygraph uses FIMO to perform motif scanning on all provided sequences with either the JASPAR database or a set of provided motifs. Based on the FIMO results, it computes frequency of each motif, frequency of pairs of motifs, and the relative orientation and spacing between pairs of motifs. Embedding metrics (described below) can be computed for each group of sequences based on their motif frequency. Statistical tests (described below) are used to compare the distributions of these metrics across groups, resulting in statistics for differential motif abundance, differential positioning, differential orientation, and differential spacing. Polygraph uses NMF to perform factor analysis on the motif frequency matrix. Finally, one issue with motif analysis using the full JASPAR database is that this may include motifs for TFs that are not relevant to the tissue or cell type being analyzed. Polygraph allows users to filter motifs based on tissue-specific expression of the corresponding TF in GTEx [31]. Also, given a predictive model, Polygraph can perform In Silico Mutagenesis (ISM) scoring for each base in each input sequence, and filter motifs that have a high importance score indicating their potential relevance in the system.

#### Predictive modeling analysis

Polygraph facilitates predictive modeling analysis through Enformer [9], Nucleotide Transformer [32], our pre-trained yeast and human models, or custom user-provided PyTorch models. The package allows for generating model predictions, producing model embeddings, and calculating cell-type specificity. Embedding metrics (described below) can be computed for each group of sequences based on their model embedding.

#### Guided evolution

The Polygraph evolve module enables custom sequence design. The evolve function performs iterative directed evolution on DNA sequences, in which all possible single-base mutations to the input sequence are explored in each iteration and the one with the best model prediction is selected for the next iteration. However, Polygraph offers users the option to simultaneously optimize for both predicted activity based on a provided model and similarity to a set of reference native DNA sequences by embedding the sequences and calculating euclidean distances in the embedding space. The function allows the user to control the relative importance of activity versus similarity through a weighting parameter. Resulting sequences have high predicted activity while maintaining similarity to a native sequence population.

#### Language models

Polygraph supports computing sequence likelihood via the hyenaDNA model [27]. Embedding metrics, which can be computed using either k-mer, motif, or model embeddings include:

1. **Sequence Diversity:** Calculated as the within-group KNN distance, representing the mean Euclidean distance of each sequence to its k nearest neighbors within the same group.
2. **Distance to Reference:** Measures the Euclidean distance of each sequence to its nearest neighbor in the reference group, within the embedding space.
3. **1-NN Statistic:** Computes the fraction of synthetic sequences in each group that have a reference sequence as their nearest neighbor based on Euclidean distance in the sequence embeddings. It can also be applied to calculate the fraction of nearest neighbors in any group.
4. **Differential Analysis:** Conducts a Wilcoxon rank-sum test on sequence embeddings to identify features enriched or depleted relative to the reference group.
5. **Classifier Performance:** Assesses the ability of a support vector machine (SVM) classifier to differentiate between each group of synthetic sequences and the reference group.

Polygraph supports statistical analyses to compare any of these metrics across groups:

1. **Groupwise Fisher’s exact test**: for comparing proportions between each non-reference group and the reference group.
2. **Groupwise Mann-Whitney U test**: for comparing mean values between each non-reference group and the reference group.
3. **Kruskal-Wallis test followed by Dunn’s post-hoc test**: for comparing mean values across all groups.

### Yeast promoter dataset and model

Yeast gigantically parallel reporter assay (GPRA) data as well as expression measurements for native promoters were obtained from Vaishnav et al.[29]. A sequence-to-expression model was trained using a convolutional architecture on 3,922 80bp sequences of native yeast promoters whose expression was measured in complex medium. The model consisted of 4 convolutional layers each with 64 channels and ReLU activation. The first convolutional layer had kernel size=15 and subsequent layers had kernel size=3. The convolutional layers were followed by average pooling across the sequence length. Finally, a 1x1 convolutional layer was used to combine the output of the 64 channels into the predicted expression value. 3,500 randomly selected promoters were used for training while 422 were held out each for validation and testing. The model was trained for 10 epochs with Poisson loss using the Adam optimizer with learning rate=1e-4 and batch size 16. This model was used as an oracle for sequence design.

To generate sequences using directed evolution, we initialized 50 random starting sequences and iterated through 10 rounds of evolution using the yeast model as an oracle to increase expression. 50 Ledidi[13] optimized sequences were generated similarly with the following parameters: max_iter=500, l=100, lr=1e-2. 50 sequences were generated through 25 iterations of the custom guided evolution function, with the native high sequences serving as the reference population.

### Yeast embedding model

We considered approximately 7.5 million 80 bp long randomly generated yeast promoters whose expression was measured in both complex and defined media from Vaishnav et al. 2022[29]. 1,000,000 randomly selected sequences were held out for training and validation. We trained a sequence-to-expression model to predict expression in both media. The model consisted of 4 convolutional layers with 512 channels each and ReLU activation. All convolutional layers after the first were followed by max pooling with width 2. The output of the final convolutional layer was flattened and passed through a dense layer with 32 nodes and ReLU activation to produce two outputs.

The output from the final convolutional layer of this model was used to embed native and synthetic sequences. The trained model can be downloaded and used as part of the Polygraph package.

### Human enhancer dataset and model

Gosai et al.[11] generated a dataset of 776,474 200-nucleotide sequences paired with gene expression, as measured by a massively parallel reporter assay (MPRA), and engineered cell-type specific candidate sequences using simulated annealing, FastSeqProp[12], and AdaLead[14]. To compare the performance of these methods in one cell type, we selected the top 2,000 synthetic sequences with the highest MinGap in K562 cells, defined by Gosai et al. as sequences with the highest difference in predicted activity in K562 and the max of HepG2 and SK-N-SH. The ‘native’ group was selected from the top 2,000 endogenous sequences overlapping H3K27ac peaks with the highest MinGap in K562 (referred to as ‘DHS-natural’).

We finetuned a pretrained Enformer model[9] using the Enformer PyTorch implementation at https://github.com/lucidrains/enformer-pytorch on the single-cell atlas of chromatin accessibility in the human genome[30] to predict binarized cell type-specific pseudobulk accessibility and embed sequences (**Fig. 3d-e, Fig S3 and S4**). Sequences from chromosomes 7 and 13 were held out for validation and testing, respectively. We sliced the first residual block and left out the next six blocks, used “target_length=-1” argument to disable cropping and added a 1D average pooling to collapse the sequence dimensionality. Input length was 200bp. Binary signal and cell type annotations were obtained from publicly available files provided by the authors (http://catlas.org/catlas_downloads/humantissues/cCRE_by_cell_type/). Cell types with less than 3% positive labels were discarded which reduced the number of cell types from 222 to 203.

## Declarations

### Availability of data and materials

Project name: Polygraph Project home page: https://genentech.github.io/polygraph, https://github.com/Genentech/polygraph

Archived version: Model weights and preprocessed datasets can be found at the following DOI:

10.5281/zenodo.10988854.

Operating system(s): Platform independent

Programming language: Python

Other requirements: Polygraph requires Python >= v3.8. A GPU is not required but supports faster model-based embeddings and predictions.

License: MIT

Any restrictions to use by non-academics: None

## Author Contributions

AG, AL, and GE formulated the project. AL developed the polygraph codebase, with support from AG and LG. AL wrote code tutorials, with support from LG. AL integrated and trained models, with support from GE. LG performed analysis on the human and yeast datasets, with support from AL and GE. LG drafted the figures and manuscript. TB provided supervision and mentorship. All authors revised the manuscript.

## Acknowledgments

We thank David Garfield and Oriol Fornes for helpful discussion and feedback.

## Main Figures

## Supplementary Information

**Supplementary Table 1**: *Yeast promoter evaluation metrics (attached)*

**Supplementary Table 2**: *Yeast top differential k-mers*

**Supplementary Table 3**: *Yeast motif counts (attached)*

**Supplementary Table 4**: *Human enhancer evaluation metrics (attached)*

**Supplementary Table 5**: *Human motif counts (attached)*

**Supplementary Figure 1**: *Yeast model performance*.

**Supplementary Figure 2**: *NMF factor loadings*.

**Supplementary Figure 3**: *CATlas model performance*.

**Supplementary Figure 4**: *Chromatin accessibility predictions*.

## Supplementary Tables

**Supplementary Table 2:**
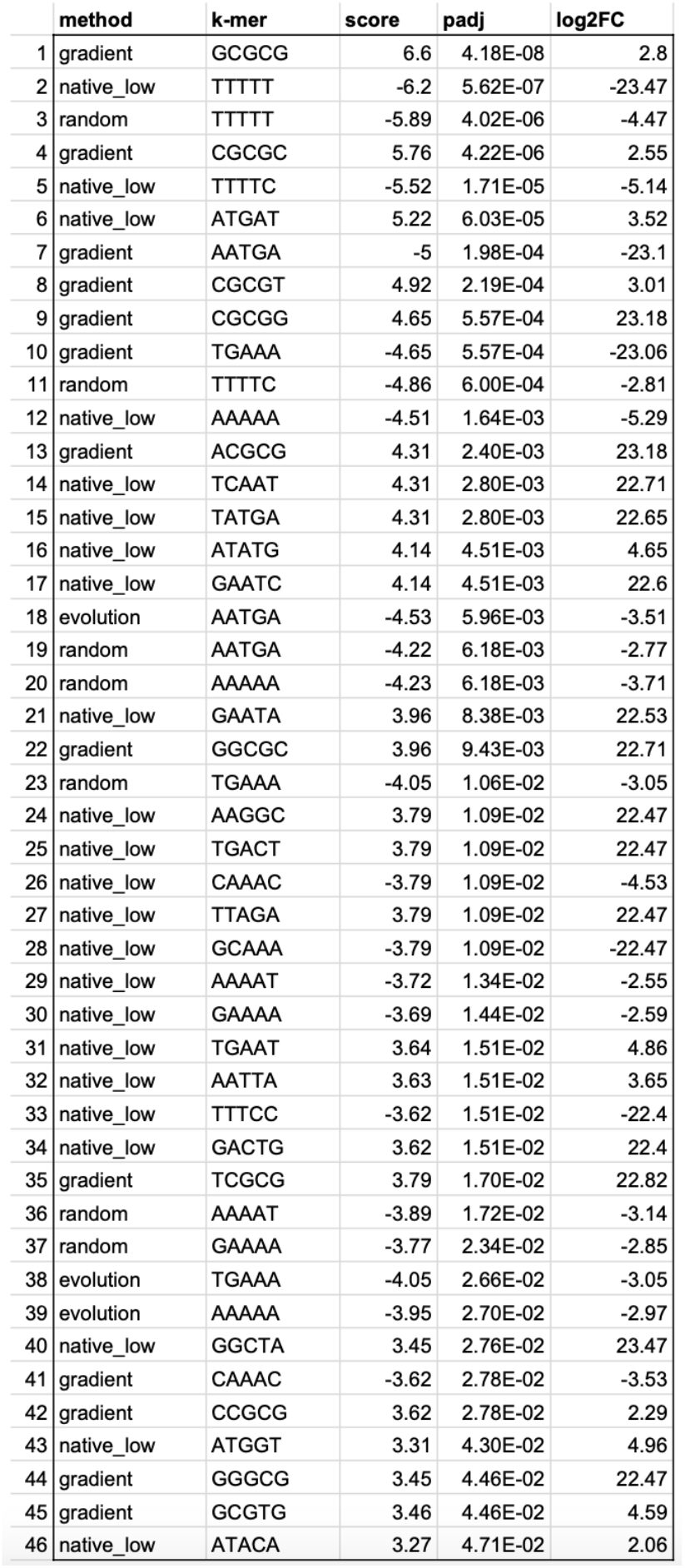
Yeast top differential k-mers. Top differential k-mers compared to native_high group (p-adj < 0.05).

## Supplementary Figures

**Supplementary Figure 1:**
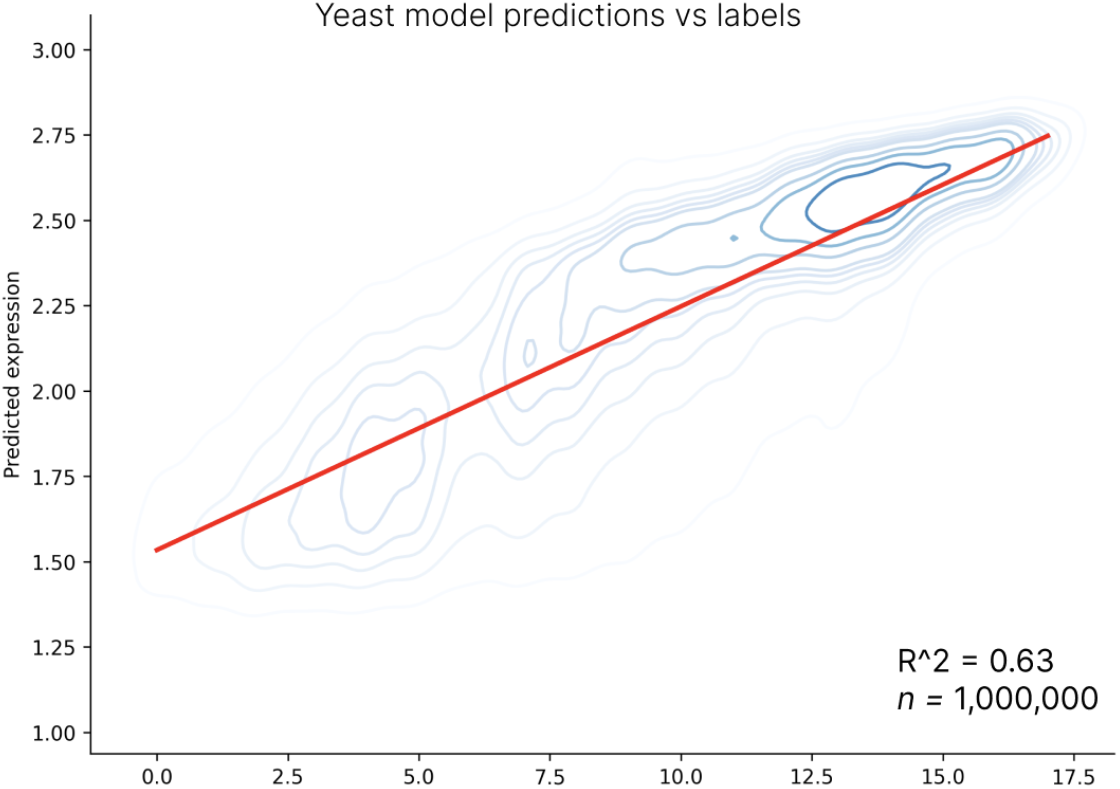
Yeast model performance. Performance of model train to predict yeast expression from random sequences on held-out dataset.

**Supplementary Figure 2:**
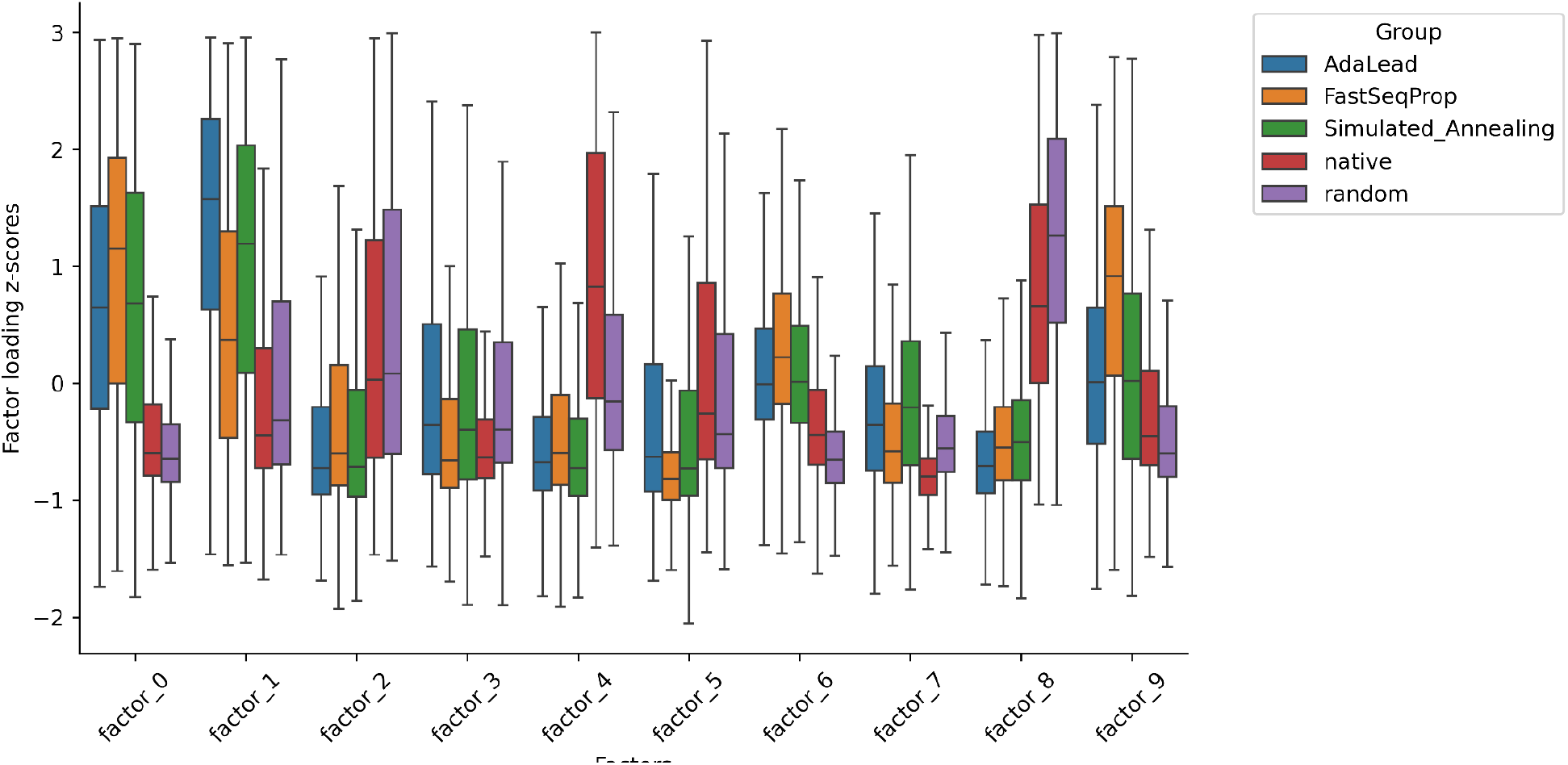
NMF factor loadings. Z-score normalized factor loadings from NMF on a motif count matrix across native, random, and synthetic human enhancer sequences.

**Supplementary Figure 3:**
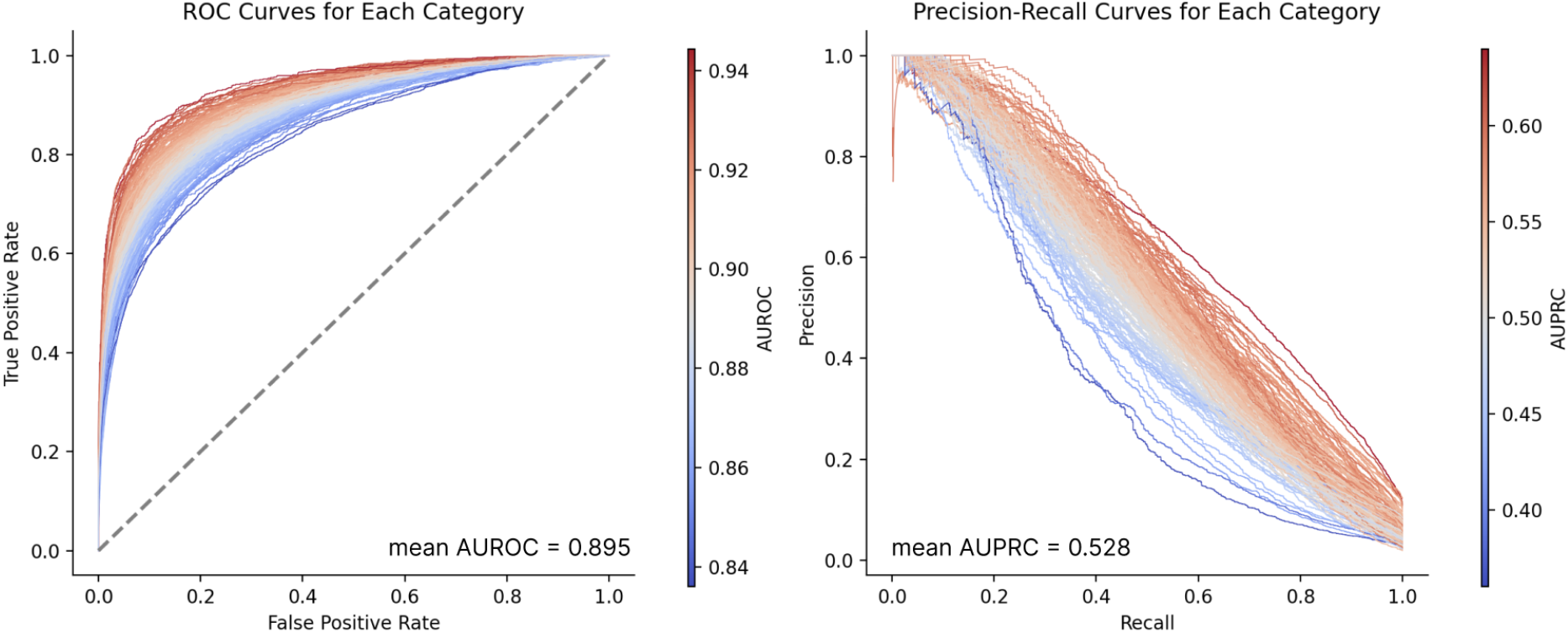
CATlas model performance. Per cell-type AUROC (left) and AUPRC (right) of a pretrained Enformer model trained to predict chromatin accessibility from sequence.

**Supplementary Figure 4:**
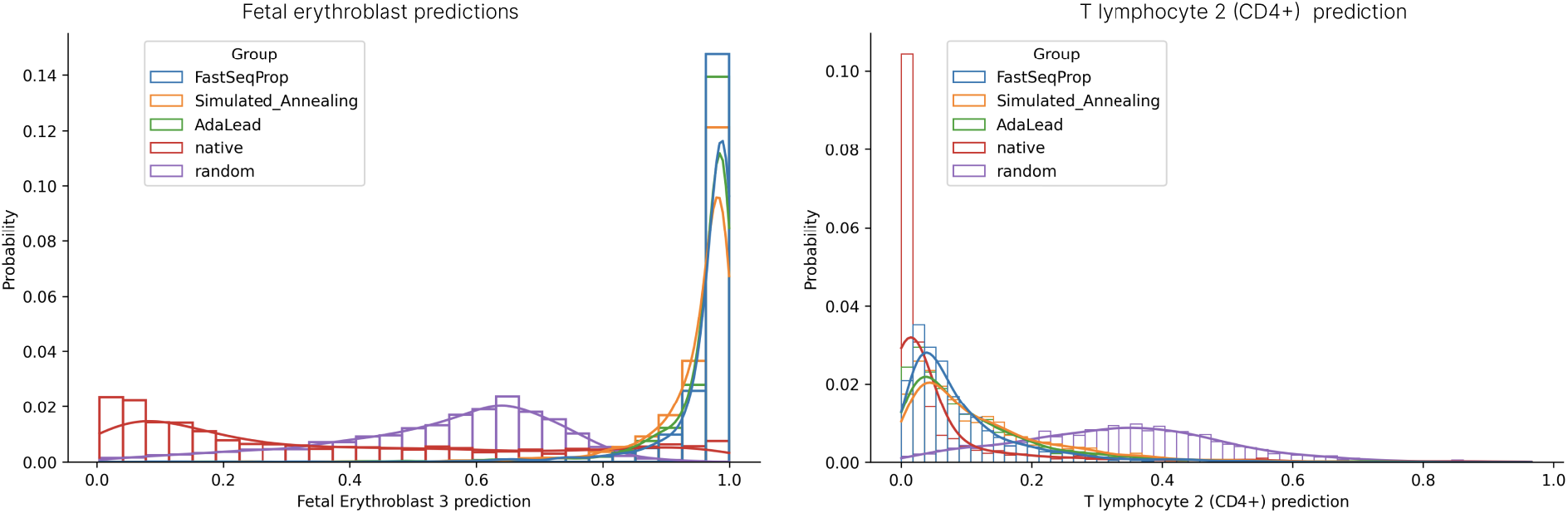
Chromatin accessibility predictions. Fetal erythroblast (left) and T lymphocyte 2 (right) ATAC-seq predictions on native, random, and synthetic K562-specific enhancer sequences.

