## Supplemental Tables for "Polygraph: A Software Framework for the Systematic Assessment of Synthetic Regulatory DNA Elements": ST1_yeast_evaluation_metrics.pdf

| | *evolution* | *gradient* | *guided evolution* | *native (high)* | *native (low)* | *random* |
|---|---|---|---|---|---|---|
| *n* | 50 | 50 | 50 | 50 | 50 | 50 |
| GC content (%) | 0.53+=0.0605 | 0.589+=0.0625 | 0.473+=0.077 | 0.391+=0.076 | 0.381+=0.0711 | 0.499+=0.0549 |
| edit distance to native high (bp) | 41.5+=1.72 | 41.7+=1.72 | 39.7+=2.19 | 0+=0 | 40.2+=1.96 | 41.6+=1.74 |
| k-mer KNN distance | 0.111+=0.0057 | 0.0929+=0.00574 | 0.0806+=0.0182 | 0.0919+=0.0263 | 0.078+=0.0271 | 0.107+=0.00637 |
| k-mer distance to reference | 0.106+=0.0115 | 0.105+=0.0101 | 0.0549+=0.00969 | 0.0221+=0.0275 | 0.127+=0.0197 | 0.108+=0.00948 |
| k-mer, fraction of sequences in 1-NN group | 0.26 | 0.02 | 0.26 | | 0 | 0.1 |
| kmer SVM accuracy | 0.97 | 0.99 | 0.88 | | 0.99 | 0.99 |
| motif KNN Distance | 0.115+=0.0195 | 0.133+=0.0136 | 0.0975+=0.0117 | 0.0746+=0.0199 | 0.0484+=0.0181 | 0.0813+=0.0141 |
| motif distance to closest reference | 0.108+=0.0294 | 0.143+=0.0323 | 0.0824+=0.0156 | 0.0292+=0.0249 | 0.0676+=0.0117 | 0.0805+=0.0149 |
| motif, fraction of sequences in 1-NN group | 0.32 | 0.2 | 0.58 | | 0 | 0.04 |
| motif SVM accuracy | 0.85 | 0.85 | 0.83 | | 0.99 | 0.93 |
| model predictions | 2.68+=0.0834 | 2.73+=0.0642 | 2.57+=0.221 | 2.76+=0.0782 | 1.54+=0.187 | 2.19+=0.403 |
| MinGap | 0.0303+=0.0108 | 0.0301+=0.01 | 0.0311+=0.0222 | 0.0519+=0.0428 | 0.166+=0.0601 | 0.0569+=0.0454 |
| model KNN distance | 2.17+=0.431 | 2.04+=0.494 | 2.61+=1.08 | 3.17+=1.39 | 1.29+=0.485 | 2.13+=0.39 |
| model distance to reference | 2.43+=0.59 | 3.33+=1.09 | 2.44+=0.632 | 0.835+=0.852 | 4.07+=0.467 | 2.73+=0.52 |
| model, fraction of sequences in 1-NN group | 0.04 | 0.02 | 0.16 | | 0 | 0.02 |
| model, SVM accuracy | 1 | 1 | 1 | | 1 | 1 |
