## Supplemental Tables for "Polygraph: A Software Framework for the Systematic Assessment of Synthetic Regulatory DNA Elements": ST4_human_evaluaton_metrics.pdf

***Human evaluation metrics***

| | *native* | *random* | *AdaLead* | *FastSeqProp* | *Simulated Annealing* |
|---|---|---|---|---|---|
| *n* | 2000 | 2000 | 2000 | 2000 | 2000 |
| GC content (%) | 0.492432 ± 0.06556 | 0.49908 ± 0.034127 | 0.498113 ± 0.023456 | 0.504595 ± 0.027641 | 0.50043 ± 0.026049 |
| edit distance to native high (bp) | | 1.628602 | 95.659 ± 1.712362 | 96.1005 ± 1.68755 | 96.0335 ± 1.649769 |
| k-mer KNN distance | 0.0341+=0.0033 | 0.0274+=0.0021 | 0.0372+=0.00227 | 0.0342+=0.00218 | 0.0357+=0.00227 |
| k-mer distance to reference | 0.0307+=0.00325 | 0.0309+=0.00229 | 0.0426+=0.00374 | 0.0402+=0.00461 | 0.0398+=0.00353 |
| k-mer, fraction of sequences in 1-NN group | | 1.20E-02 | 4.00E-03 | 1.95E-02 | 9.50E-03 |
| kmer SVM accuracy | | 0.98 | 1 | 0.99 | 1 |
| motif KNN Distance | 0.0802+=0.0217 | 0.0608+=0.0122 | 0.0766+=0.0107 | 0.0735+=0.0116 | .0747+=0.0107 |
| motif distance to closest reference | 0.07+=0.0173 | 0.0623+=0.0112 | 0.0876+=0.0151 | 0.0875+=0.0194 | 0.083+=0.0146 |
| motif, fraction of sequences in 1-NN group | | 0.08 | 0.02 | 0.06 | 0.03 |
| motif SVM accuracy | | 0.97 | 0.99 | 0.98 | 0.99 |
| model predictions (fetal erythroblast) | -0.79846 ± 2.045501 | 0.162757 ± 0.890951 | 3.81648 ± 1.297389 | 4.059606 ± 1.37303 | 3.521847 ± 1.241686 |
| model KNN distance | 0.138077 ± 0.280509 | 0.114764 ± 0.351651 | 0.168709 ± 0.220516 | 0.175887 ± 0.19118 | 0.184924 ± 0.354068 |
| 1 NN distance | 0.033517 ± 0.023881 | 0.029103 ± 0.018575 | 0.032111 ± 0.019595 | 0.031111 ± 0.019917 | 0.032802 ± 0.020755 |
| model distance to reference | 0.043535 ± 0.130434 | 0.242532 ± 0.160609 | 0.936939 ± 0.659033 | 1.081314 ± 0.681797 | 0.859345 ± 0.633261 |
| log likelihood | -258.871729 ± 24.148949 | -288.94453 ± 4.027626 | -287.501586 ± 4.216266 | -288.569882 ± 3.905383 | -288.72723 ± 4.01331 |
